# Amphibian and reptile dataset across different land-use types in Guinea-Bissau, West Africa

**DOI:** 10.1101/2024.11.21.624282

**Authors:** Francisco dos Reis-Silva, Fernanda Alves-Martins, Javier Martinez-Arribas, Cristian Pizzigalli, Sambu Seck, Ana Rainho, Ricardo Rocha, Ana Filipa Palmeirim

**Affiliations:** Global Change and Conservation Research Group, Faculty of Biological and Environmental Sciences, University of Helsinki, Viikinkaari 1 P.O. Box 65 00014, Finland; CIBIO, Centro de Investigação em Biodiversidade e Recursos Genéticos, InBIO Laboratório Associado, Campus de Vairão, Universidade do Porto, 4485-661 Vairão, Portugal; BIOPOLIS Program in Genomics, Biodiversity and Land Planning, CIBIO, Campus de Vairão, 4485-661 Vairão, Portugal; Federação KAFO, Guiné-Bissau CP 1186, Centro Camponês de Djalicunda, sector de Mansaba, região de Oio, Guinea-Bissau; Centre for Ecology, Evolution and Environmental Changes & CHANGE - Global Change and Sustainability Institute. Departamento de Biologia Animal, Faculdade de Ciências, Universidade de Lisboa, 1749-016 Lisboa, Portugal; Department of Biology, University of Oxford, 11a Mansfield Rd, OX1 3SZ, Oxford, UK; EcoHealth Alliance, New York City, NY, United States

**Keywords:** Amphibians, agroecosystems, Guinea-Bissau, habitat conversion, species diversity, reptiles, tropical forest, Wallacean shortfall

## Abstract

West Africa is exceptionally biodiverse, yet its wildlife remains largely understudied despite the rapid and ongoing land-use changes. Large swaths of Guinea-Bissau’s landscape were historically characterized by native forest-savanna mosaics. However, key areas of savannah habitats have been converted to rice agroecosystems, and forests are being transformed into cashew monocultures at unprecedented rates. Amphibians and reptiles (Testudines and Squamata) comprise some of the most threatened species by human-induced habitat change, and yet are not as studied as other vertebrate terrestrial taxa. Here, we provide two comprehensive datasets on amphibians and reptiles (classes Testudines and Squamata) from northern Guinea-Bissau: (1) a standardized survey dataset (encompassing sampling events and occurrences) in forest fragments, cashew orchards, and rice paddies, and (2) an opportunistic dataset reporting occurrences across the entire study area. Standardized surveys were carried across 21 sampling sites, seven in each habitat type, while opportunistic surveys include all other records. For standardized surveys, a total of 703 amphibian and 265 reptile (class Squamata) encounters are reported, corresponding to nine and 13 taxa, respectively. Opportunistically, we report 62 amphibian and 93 reptile encounters, corresponding to 10 amphibian taxa, 25 Squamata taxa, and two turtles (class Testudines).

**New information:** Based on 126 sampling hours of both diurnal and nocturnal standardized surveys, in addition to opportunistic surveys, these datasets comprise the first overview for amphibians and reptiles in mainland Guinea-Bissau across two seasons and different habitat types. Each of the 968 standardized and 155 opportunistic occurrences corresponds to a genus or species and is accompanied by geographic coordinates, a timestamp, and, for standardized data, the land-use type. The datasets fill the distribution gaps in Guinea-Bissau of at least three species, including the frog *Hildebrandtia ornata*, the skink *Trachylepis keroanensis* and the snake *Dendroaspis polylepis* – and includes the rediscovery of the lizard *Latastia ornata* in Guinea-Bissau. Before this work, the *L. ornata* was only known from the 1938 holotype in Bafatá (ca. 60 km away from the study area) and, in 2023, from Guinea-Conakry (ca. 700 km away from type specimen location).

## Introduction

West Africa is a major biodiversity hotspot, with a high number of endemic species (Myers et al. 2000). The region has faced substantial habitat loss and degradation (Lewin et al. 2016), which is expected to continue (Powers and Jetz 2019). Yet, West Africa has been subject to very few ecological studies compared to other biodiversity hotspots, such as the Neotropics (Gardner et al. 2009, Gibson et al. 2011, Newbold et al. 2020).

Guinea-Bissau has been covered by native forest-savanna mosaics (Catarino et al. 2008), but its long history of agriculture has changed the landscape overtime (Temudo and Abrantes 2013). Rice (*Oryza L*.) has traditionally been cultivated for domestic use (Temudo and Abrantes 2013) and, together with groundnuts, comprised the core of the agricultural land in the country until the 20^th^ century (Catarino et al. 2015). After the 1940’s, cashew trees (*Anacardium occidentale* L.) – native to Northeast Brazil – started being systematically planted across the country (Temudo and Abrantes 2014). This global agricultural commodity (Rege and Lee 2023) has replaced most other forms of land use in Guinea-Bissau, especially since the 1980’s (Temudo and Abrantes 2013). Today, agriculture is still the main source of livelihood in the country, with cashew nuts comprising the only cash crop for the economy of Guinea-Bissau (Temudo and Abrantes 2013), accounting for 90% of all exports (FAO 2021). The once highly complex bio-cultural landscapes in Guinea-Bissau are now threatened by the quick expansion of cashew orchards, which are homogenizing the landscape (Catarino et al. 2015).

Amphibians and reptiles (classes Testudines and Squamata) are among the most threatened vertebrates (Cox et al. 2022), but their responses to anthropogenic pressure are less studied than that of other taxa (e.g., invertebrates and birds; Newbold et al. 2014) and there is a strong geographical bias in the available literature, with efforts skewed toward temperate regions and the Neotropics (Guedes et al. 2023, Tan et al. 2023). This has led to species such as the West African lizard *Latastia ornata* Monard, 1940 being only known from one type locality specimen for over 80 years (Meiri et al. 2018, Pauwels et al. 2023), or the medically significant black mamba *Dendroaspis polylepis* Günther, 1864 having only a few scattered observations in West Africa when, in fact, its distribution is suspected to be continuous in the region (Chippaux and Jackson 2019). The Wallacean shortfall, particularly evident in West African herpetofauna, reflects this geographic bias in species distribution data, which has large implications for species’ conservation (Hortal et al. 2015) and human well-being, as this lack of information, as seen with venomous snakes, contributes to a higher incidence of untreated snakebites (WHO 2023). Despite the scarcity of scientific studies, amphibians and reptiles are deeply embedded in the region’s biocultural heritage (e.g., notwithstanding considerable levels of disliked towards snakes, Bissau-Guinean farmers often perceive snakes as protectors of the village and signs of a good harvest; Chaves et al., in press).

To help filling in the knowledge gap in amphibian and reptile distribution in West Africa, we provide two herpetofauna datasets resulting from standardized and opportunistic surveys across the study area. The habitat mosaic, encompassing forest remnants, cashew orchards and rice paddies, was specifically chosen to detect a wide range of species associated with both open- and closed-habitats. Furthermore, it informs us of species that can be found in the expanding cashew orchards. Unconventionally, the surveys were carried outside protected areas, which contributes further to overcoming the Wallacean Shortfall.

## Sampling methods

### Study area

This study took place in northern Guinea-Bissau, Oio Province, in the surroundings of Djalicunda (12°19’49.82”N, 15°10’57.55”W, Figure 1). The region’s landscape consists of scattered small *tabancas* (villages) surrounded by secondary forest and large areas of extensive smallholder agriculture. The semi-natural and agricultural areas create mosaics of mostly forest remnants, cashew orchards and rice paddies. Within the region, cashew orchards are gaining prominence, leading to the clearing of some of the forest remnants (Temudo and Abrantes 2014). The area is mostly flat, below 50 m altitude, and has defined wet – from June to October – and dry – from October to June – seasons (Catarino et al. 2008). The mean temperature in the country ranges between 25.9 and 27.1 ºC, and the annual precipitation between 1200 mm in the northeast and 2600 mm in the southwest (Catarino et al. 2008). The amphibian and reptile surveys were conducted mainly across three habitat types: forest remnants, cashew orchards and rice paddies. The surveys took place in 21 study sites, seven of each habitat type.

**Figure 1.**
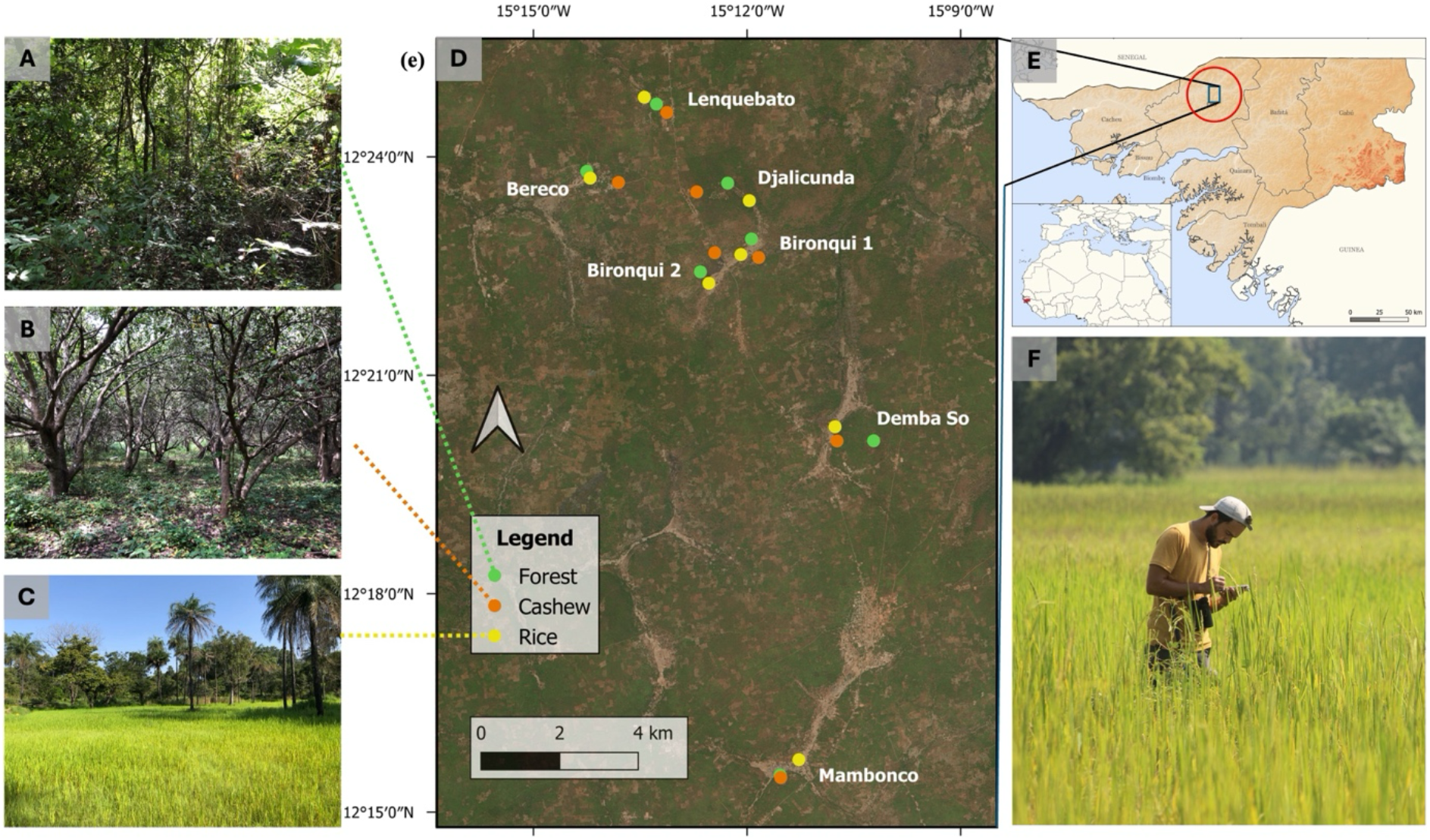
Study area and surveyed habitat types. A) Forest remnants. C) Rice paddies. D) Overview of the study area, including study sites (colored dots corresponding to the habitat type on the legend). B) Cashew orchards. E) Study area in northern Guinea-Bissau. F) Example of a survey conducted in a rice paddy. Map sources: qGIS (2023), GADM (2021) and geoBoundaries (2017). Photos: Francisco Reis-Silva.

### Surveyed habitat types

Forest remnants in the study area are classified as secondary growth, as they are either heavily degraded or represent regrowth following human intervention (Catarino et al. 2008). In the surveyed forest remnant sites, products (e.g., wood, fruit, honey) are collected by local communities, and the ground is typically covered by leaf litter, and the canopy cover is *≥*65% (dos Reis-Silva et al. in press.). Surveyed cashew orchards sites are monocultures subject to little management (i.e., no irrigation, no fertilizers). They are characterized by a dense canopy (usually *≥*80%) about 6-10 meters above the ground, and the understory is cleared once a year to facilitate cashew nut harvest (dos Reis-Silva et al. in press). Rice paddies are in topographic depressions that flood naturally between late July and November, which coincides with the plantation and harvesting of rice, respectively (Sottomayor et al. 2024). They have few scattered trees throughout, presenting an open habitat without canopy cover.

### Standardized herpetofauna surveys

Data collection took place over two field campaigns in 2022. To maximize the number of recorded species given the strong seasonality in the study area, the first field campaign occurred at the end of the dry season (June/July), and the second one at the end of the wet season (October/November). For each campaign, all sampling sites were surveyed three times during the day (starting between 09h15 and 16h45) and once at night (starting between 19h00 and 22h45), totalling eight surveys at each of the 21 sites (six day- and two night-surveys).

Herpetofauna surveys took place across 21 circular study sites of 25 m radius in time-standardized fashion (dos Reis-Silva et al. in press). Surveys were systematically conducted by one observer for 45 minutes, amounting to a total of 126 sampling hours: 94.5 h during daytime and 31.5 h during nighttime. In each survey, the study sites were thoroughly searched in a zig-zag fashion and carefully checked for herpetofauna, including underneath loose objects (e.g., dead wood, bark, leaf litter). We took note of the date and time at the beginning of each survey. For each encounter (i.e., observed individual), species and genus were registered. At times, photos were used for ID confirmation. On some occasions, no animals were detected at a study site. These zero-encounter surveys were excluded from the standardized dataset, as the lack of observations does not necessarily indicate true species absence (MacKenzie et al. 2017).

### Opportunistic herpetofauna surveys

These surveys took place throughout the study area and included all records collected outside of the standardized surveys’ locations. As such, opportunistic surveys include all amphibians and reptiles (classes Squamata and Testudines) observed while commuting to and between sampling sites and at the accommodation surroundings. Additionally, specimens found by locals, which identification we were able to confirm (e.g., road kills), were also included as opportunistic records.

### Species identification

Herpetofauna was identified visually based on morphological characters. On some occasions deemed needed and safe, animals were caught for identification (e.g., ridge-count for frogs, scale-count for reptiles). Amphibians were identified with the aid of AmphibiaWeb (AmphibiaWeb 2022) and other scientific literature (Pickersgill 2007, Auliya et al. 2012). For reptile identification, Reptile Database (Uetz et al. 2023), and the field guides Chippaux and Jackson (2019) for snakes and Trape et al. (2012) for lizards and testudines were used. Due to the lack of conclusive unique morphological characters for some species and several specimens hiding quickly, conclusive identification to species level was not always possible. Consequently, 412 observations of amphibians in the standardized dataset and 32 observations (27 amphibians and four squamates) in the opportunistic dataset were identified only at the genus level. Because the datasets only include specimens identified accurately to genus or species level, one record identified to the family Leptotyphlopidae is excluded.

## Geographic coverage

### Description

The study took place in northern Guinea-Bissau, Oio Province, in the surroundings of Djalicunda.

### Coordinates

Standardized survey: Latitude: between 12.258 and 12.414; Longitude: between -15.17 and -15.238. Opportunistic occurrences: Latitude: between 12.258 and 12.522; Longitude: between -15.169 and - 15.238.

## Taxonomic coverage

### Description

The dataset from standardized surveys includes a total of 703 amphibian and 265 squamates encounters, corresponding to nine amphibian and 13 squamate taxa (Table 1, Figure 2). In contrast, the dataset from opportunistic surveys includes 62 amphibian, three testudines and 90 squamates encounters, corresponding to 10 amphibian taxa, two testudine taxa and 25 squamate taxa (Table 2, Figures S1, S2).

**Table 1.**
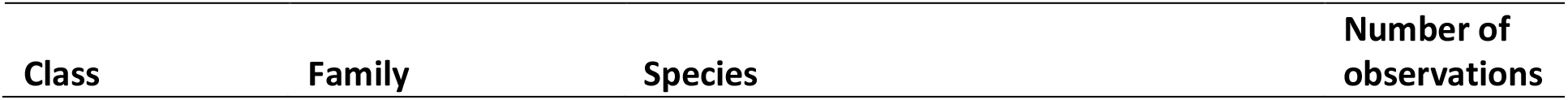
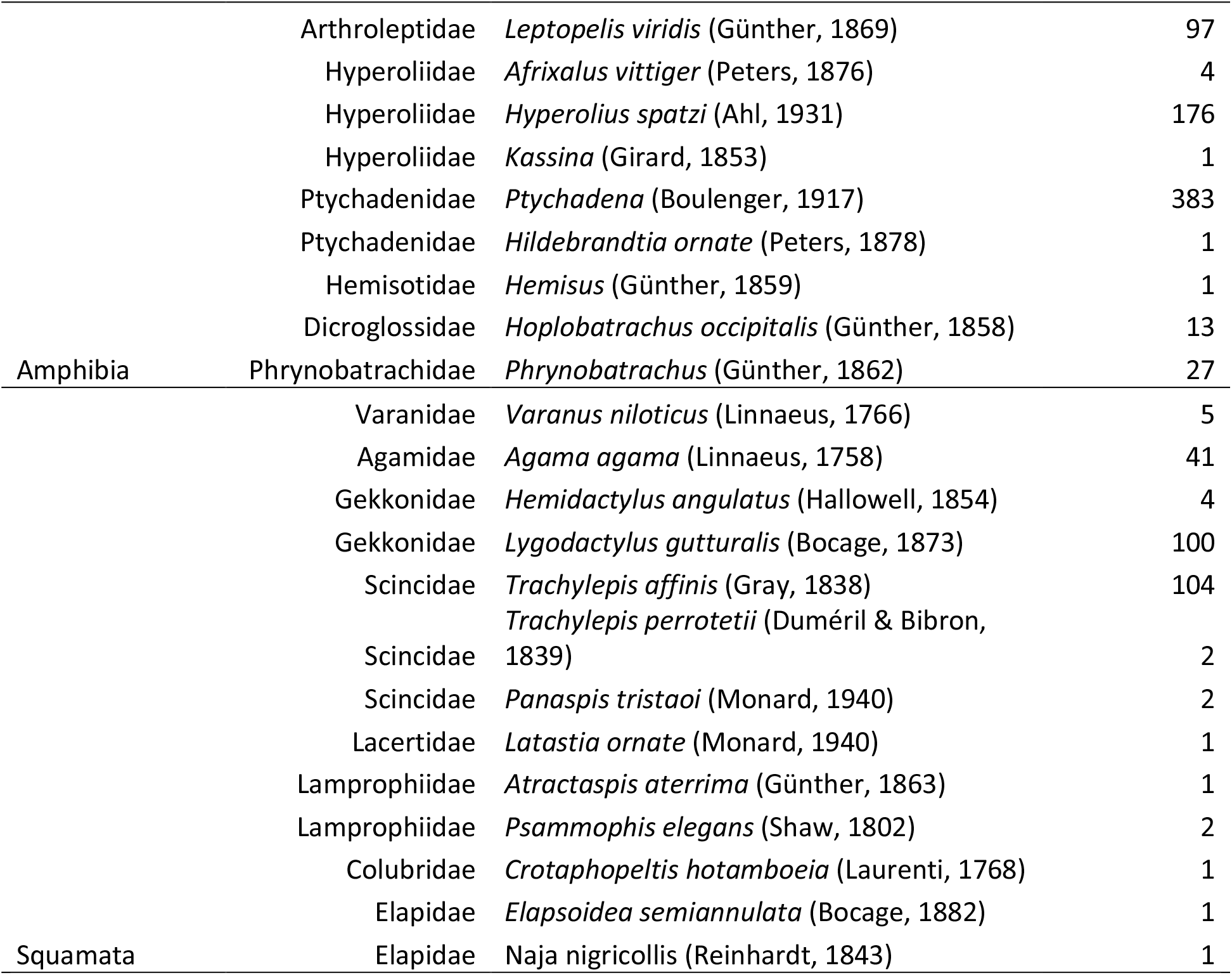
Amphibian and reptile (Squamata) observations during standardised surveys in northern Guinea-Bissau, West Africa.

**Table 2.**
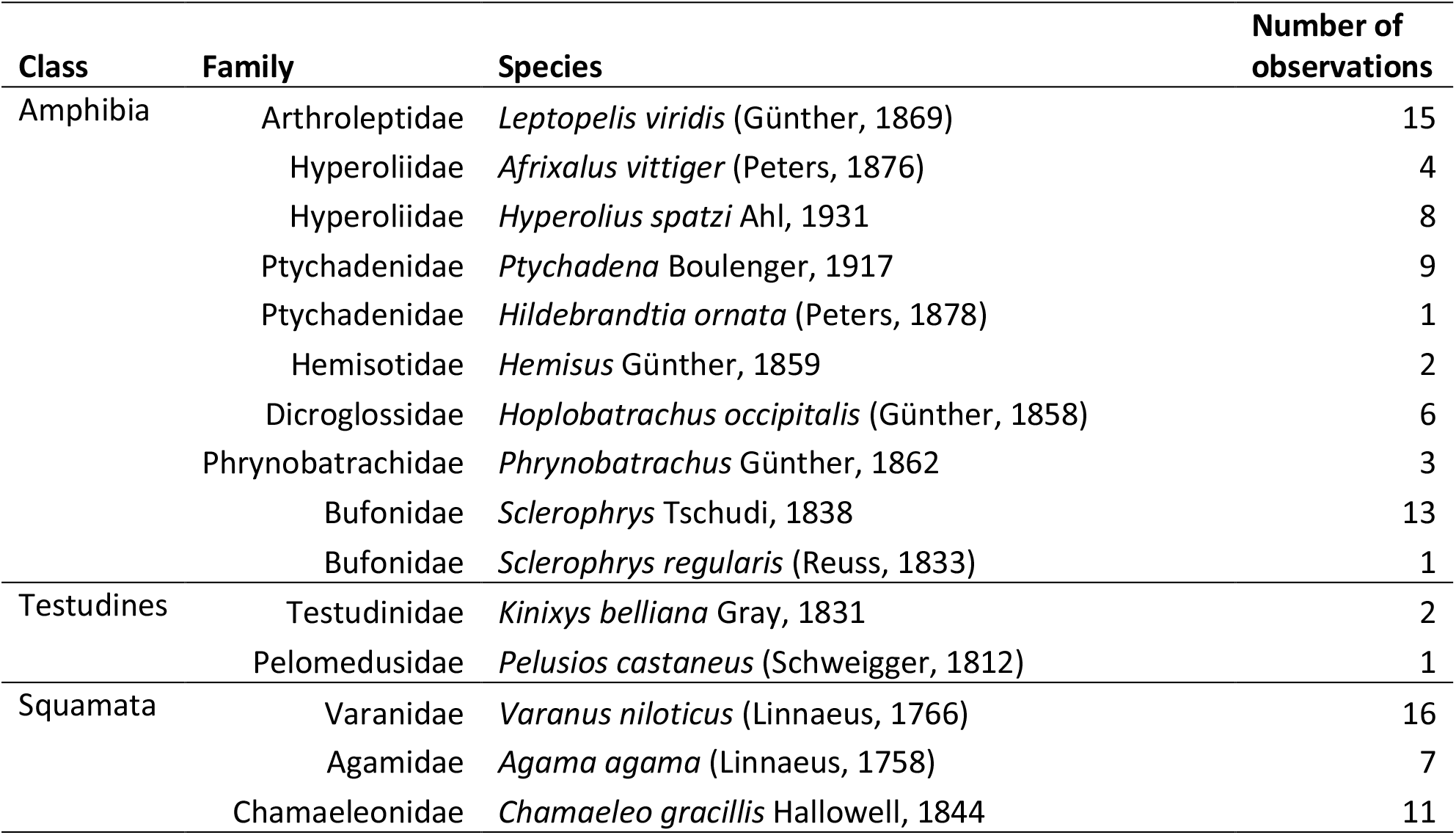
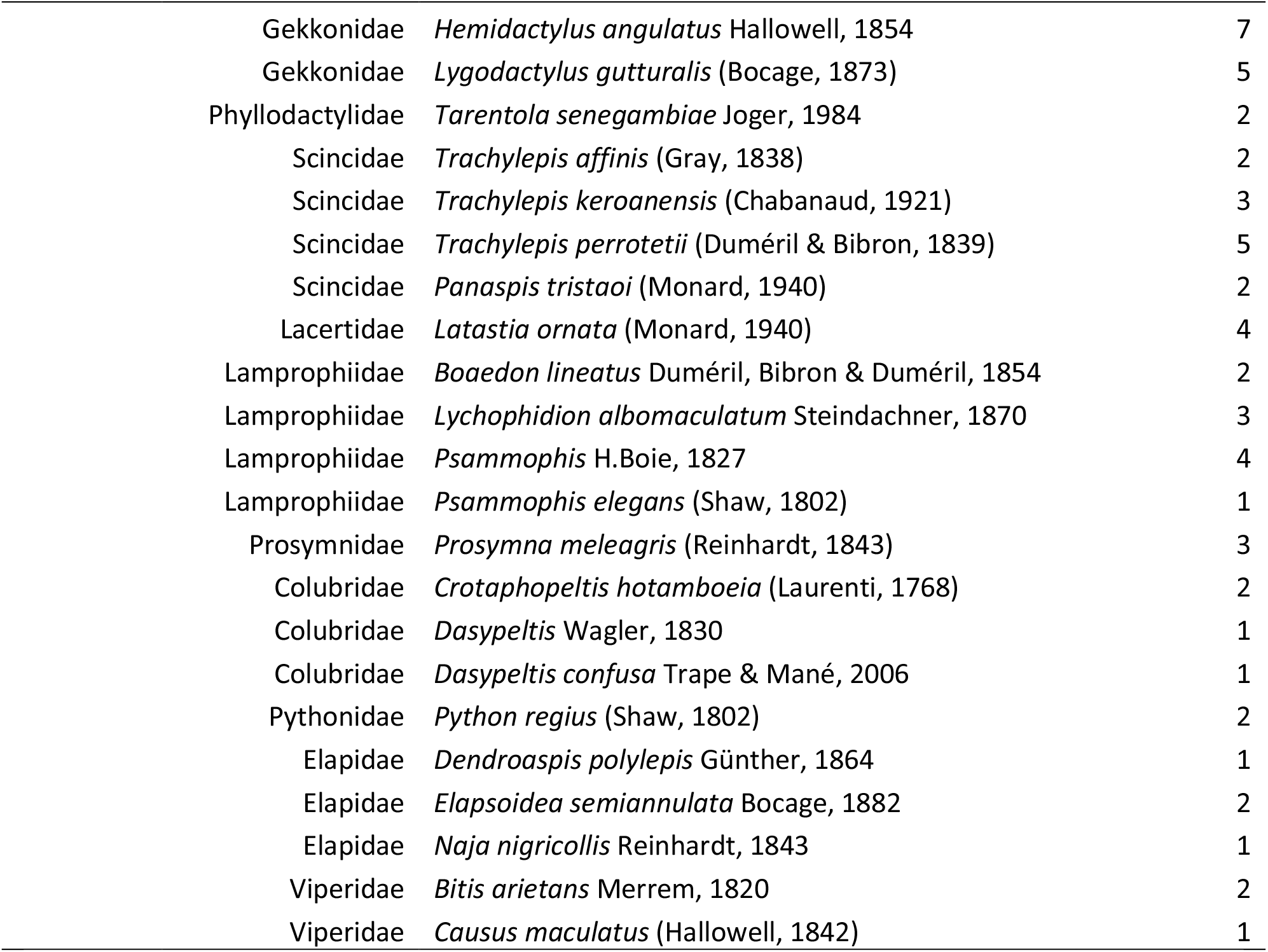
Amphibian and reptile (classes Testudines and Squamata) opportunistically detected in Guinea-Bissau, West Africa.

| Class | Family | Species | Number of observations |
| --- | --- | --- | --- |
| Amphibia | Arthroleptidae | *Leptopelis viridis* (Günther, 1869) | 15 |
|  | Hyperoliidae | *Afrixalus vittiger* (Peters, 1876) | 4 |
|  | Hyperoliidae | *Hyperolius spatzi* Ahl, 1931 | 8 |
|  | Ptychadenidae | *Ptychadena* Boulenger, 1917 | 9 |
|  | Ptychadenidae | *Hildebrandtia ornata* (Peters, 1878) | 1 |
|  | Hemisotidae | *Hemisus* Günther, 1859 | 2 |
|  | Dicroglossidae | *Hoplobatrachus occipitalis* (Günther, 1858) | 6 |
|  | Phrynobatrachidae | *Phrynobatrachus* Günther, 1862 | 3 |
|  | Bufonidae | *Sclerophrys* Tschudi, 1838 | 13 |
|  | Bufonidae | *Sclerophrys regularis* (Reuss, 1833) | 1 |
| Testudines | Testudinidae | *Kinixys belliana* Gray, 1831 | 2 |
|  | Pelomedusidae | *Pelusios castaneus* (Schweigger, 1812) | 1 |
| Squamata | Varanidae | *Varanus niloticus* (Linnaeus, 1766) | 16 |
|  | Agamidae | *Agama agama* (Linnaeus, 1758) | 7 |
|  | Chamaeleonidae | *Chamaeleo gracillis* Hallowell, 1844 | 11 |

|  |  |  |
| --- | --- | --- |
| Gekkonidae | *Hemidactylus angulatus* Hallowell, 1854 | 7 |
| Gekkonidae | *Lygodactylus gutturalis* (Bocage, 1873) | 5 |
| Phyllodactylidae | *Tarentola senegambiae* Joger, 1984 | 2 |
| Scincidae | *Trachylepis affinis* (Gray, 1838) | 2 |
| Scincidae | *Trachylepis keroanensis* (Chabanaud, 1921) | 3 |
| Scincidae | *Trachylepis perrotetii* (Duméril & Bibron, 1839) | 5 |
| Scincidae | *Panaspis tristaoi* (Monard, 1940) | 2 |
| Lacertidae | *Latastia ornata* (Monard, 1940) | 4 |
| Lamprophiidae | *Boaedon lineatus* Duméril, Bibron & Duméril, 1854 | 2 |
| Lamprophiidae | *Lychophidion albomaculatum* Steindachner, 1870 | 3 |
| Lamprophiidae | *Psammophis* H.Boie, 1827 | 4 |
| Lamprophiidae | *Psammophis elegans* (Shaw, 1802) | 1 |
| Prosymnidae | *Prosymna meleagris* (Reinhardt, 1843) | 3 |
| Colubridae | *Crotaphopeltis hotamboeia* (Laurenti, 1768) | 2 |
| Colubridae | *Dasypeltis* Wagler, 1830 | 1 |
| Colubridae | *Dasypeltis confusa* Trape & Mané, 2006 | 1 |
| Pythonidae | *Python regius* (Shaw, 1802) | 2 |
| Elapidae | *Dendroaspis polylepis* Günther, 1864 | 1 |
| Elapidae | *Elapsoidea semiannulata* Bocage, 1882 | 2 |
| Elapidae | *Naja nigricollis* Reinhardt, 1843 | 1 |
| Viperidae | *Bitis arietans* Merrem, 1820 | 2 |
| Viperidae | *Causus maculatus* (Hallowell, 1842) | 1 |

**Figure 2.**
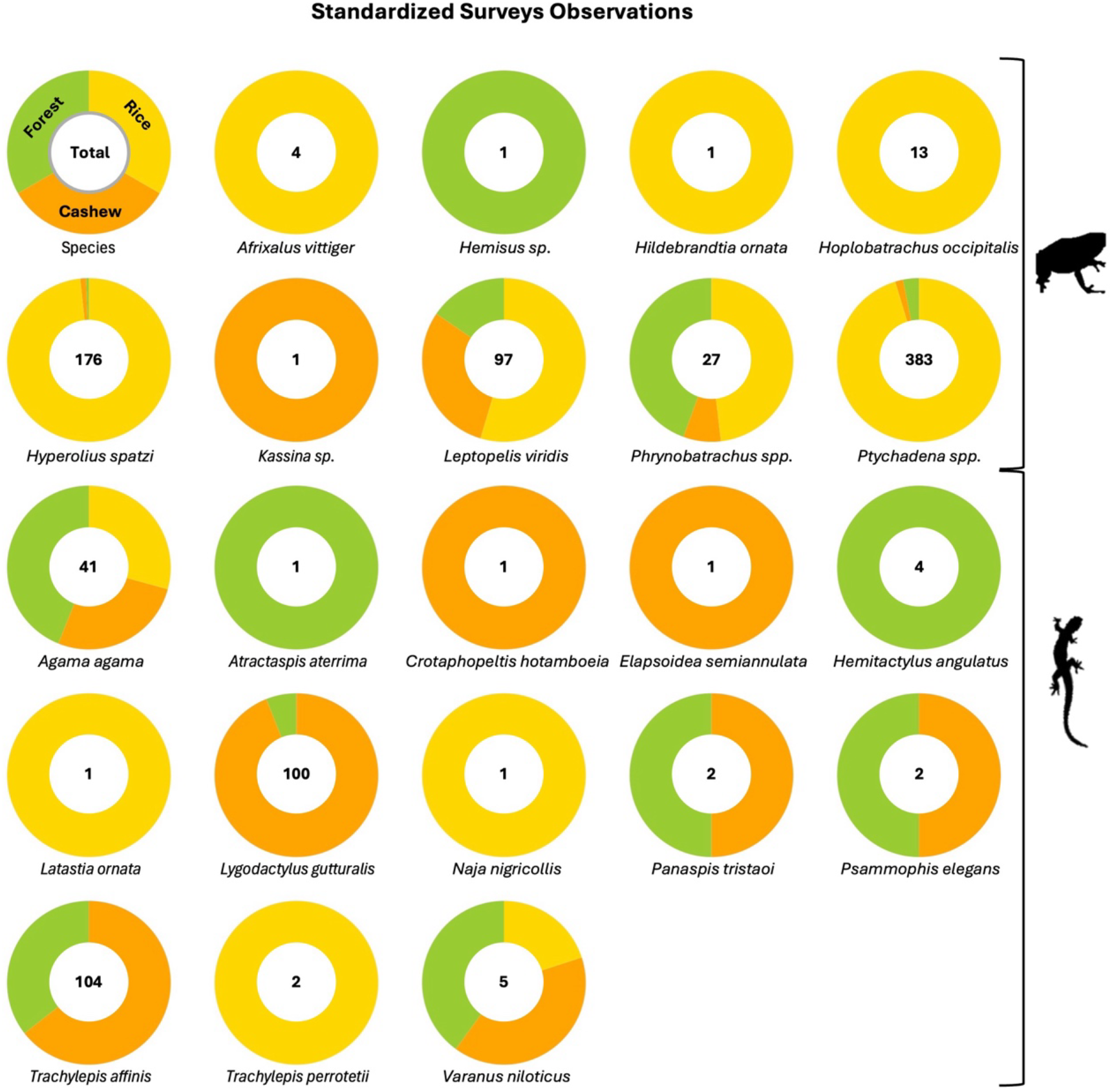
Amphibians and reptiles recorded during standardized surveys and corresponding proportion per habitat-type.

## Temporal coverage

### Data range

2022-06-18 to 2022-11-05 for DOI: 10.15468/vv9xnb (standardized survey); 2022-06-15 to 2022-11-06 for DOI: 10.15468/dwectn (opportunistic occurrences)

### Usage licence

#### Usage licence

Creative Commons Public Domain (CC-Zero)

## Data resources

### Number of data sets

2

### Data set name

Standardized survey dataset of amphibian and reptile across different land-use types in Guinea-Bissau, West Africa

### Character set

UTF-8

### Download URL

http://ipt.gbif.pt/ipt/archive.do?r=gw_herpetol_dataset

### Data format

Darwin Core Archive format

### Description

A comprehensive dataset of standardized surveys of amphibians and reptiles (Testudines and Squamata) conducted primarily across forest fragments, cashew orchards, and rice paddies in northern Guinea-Bissau is presented. Standardized surveys were conducted at 21 sampling sites, with seven sites in each habitat type. A total of 703 amphibian and 265 reptile encounters were recorded, corresponding to nine and 13 taxa, respectively.

### Data set name

Opportunistic records of amphibian and reptile across different land-use types in Guinea-Bissau, West Africa

### Character set

UTF-8

### Download URL

http://ipt.gbif.pt/ipt/archive.do?r=gw_herpetol_occurr_dataset

### Data format

Darwin Core Archive format

### Dataset description

A comprehensive dataset of opportunistic surveys of amphibians and reptiles conducted in northern Guinea-Bissau, Oio Province, in the surroundings of Djalicunda. Opportunistic surveys yielded 62 amphibian, three testudines and 90 squamates encounters, corresponding to 10 amphibian taxa, two testudine taxa and 25 squamate taxa.

## Acknowledgements

We are grateful to the workers of the NGO KAFO (Guinea-Bissau), including Ami, Belomi, Djunco, Djari, Francisco, Jara, Judite and Ioba; to the people of Djalicunda and all the *tabancas* where our fieldwork was conducted. We acknowledge financial support from *EcoPestSupression* (cE3c, Lisbon, Portugal; reference no. PTDC/ASP-AGR/0876/2020, DOI: 10.54499/ PTDC/ASP-AGR/0876/2020) and *TROPIBIO* (CIBIO, Vairão, Portugal; European Union’s Horizon 2020 research and innovation programme under grant agreement no. 854248), CP was supported by FCT-Fundação para a Ciência e Tecnologia (grant number 2020.05054.BD). We also thank Mutaro Camará and Paula Lopes for contributing with herpetofauna observations around the research station; and Amaia Gonzaga Roa, Aina Rossinyol Fernàndez and Daniel Fernández García who helped producing the maps and plots. Lastly, we thank Mar Cabeza for facilitating the project through connecting the Global Change and Conservation Group at the University of Helsinki with the Research Center for Biodiversity and Genetic Resources at the University of Porto, without which this project wouldn’t have been possible.

## Author contributions

FRS, RR, AR and AFP conceived and designed the methodology. SS settled the logistics required for the field work. FRS, RR and CP collected the data and identified the specimens. JM-A and FA-M curated the data. FRS led the writing. All authors contributed critically to the drafts and gave final approval for publication.

## References

ArcGIS [ArcMap]. (2023). Release 10.1. Redlands, CA: Environmental Systems Research Ins4tute, Inc., 2010.

AmphibiaWeb. (2022). https://amphibiaweb.org. University of California, Berkeley, CA, USA. Accessed on 15.05.2022.

Auliya, M., Wagner, P., & Böhme, W. (2012). The herpetofauna of the Bijagós archipelago, Guinea-Bissau (West Africa) and a first country-wide checklist. Bonn zoological Bulletin, 61(2), 255–281.

Catarino, L., Martins, E. S., Basto, M. F. P., & Diniz, M. A. (2008). An Annotated Checklist of the Vascular Flora of Guinea-Bissau (West Africa). Blumea - Biodiversity, Evolution and Biogeography of Plants, 53(1), 1–222. 10.3767/000651908X608179.

Catarino, L., Menezes, Y., & Sardinha, R. (2015). Cashew cultivation in Guinea-Bissau – risks and challenges of the success of a cash crop. Scientia Agricola, 72(5), 459–467. 10.1590/0103-9016-2014-0369.

Chaves, P. A. P., Schaafsma, M., Dabo, D., Lomba, J. Z., Mane, F., Lima, R. F., Parlmeirim, J. M., Rocha, R., Seck, S., Biaj, J., Timóteo, S., Meyer, C. F. J., Rainho, A. (in press). Friend or Foe? A`tudes of Rice Farmers towards Wild Animals in West Africa. Ecology and Society.

Cox, N., Young, B. E., Bowles, P., Fernandez, M., Marin, J., Rapacciuolo, G., … & Xie, Y. 471 (2022). A global reptile assessment highlights shared conservation needs of 472 tetrapods. Nature, 605(7909), 285–290. 10.1038/s41586-022-04664-7.

Chippaux, J.-P., & Jackson, K. (2019). Snakes of Central and Western Africa. Johns Hopkins University Press.

Database of Global Administra4ve Areas (GADM). (2021). Africa Boundaries. Database of Global Administrative Areas (GADM). Retrieved from https://hub.arcgis.com/datasets/07610d73964e4d39ab62c4245d548625_0/about.

dos Reis Silva, F., Pizzigalli, C., Seck, S., Cabeza, M., Rainho, A., Rocha, R., Palmeirim, A. F. (in press). Biotropica.

FAO. (2021). https://www.fao.org/countryprofiles/news-archive/detail-news/en/c/1471318/. Accessed on 01.05.2023.

Fulgence, T. R., Martin, D. A., Randriamanantena, R., Botra, R., Befidimanana, E., Osen, K., Wurz, A., Krev, H., Andrianarimisa, A., & Ratsoavina, F. M. (2021). Differential responses of amphibians and reptiles to land-use change in the biodiversity hotspot of north-eastern Madagascar. Animal Conservation, acv.12760. 10.1111/acv.12760.

Gardner, T. A., Barlow, J., Chazdon, R., Ewers, R. M., Harvey, C. A., Peres, C. A., & Sodhi, N. S. (2009). Prospects for tropical forest biodiversity in a human-modified world. Ecology Leiers, 12(6), 561–582. 10.1111/j.1461-0248.2009.01294.x.

geoBoundaries - Global Database of Political Administrative Boundaries. (2017). Guinea-Bissau - Subnational Administrative Boundaries. geoBoundaries-GNB-ADM1-all.zip.. Retrieved from https://data.humdata.org/dataset/geoboundaries-admin-boundaries-for-guinea-bissau.

Gibson, L., Lee, T. M., Koh, L. P., Brook, B. W., Gardner, T. A., Barlow, J., Peres, C. A., Bradshaw, C. J. A., Laurance, W. F., Lovejoy, T. E., & Sodhi, N. S. (2011). Primary forests are irreplaceable for sustaining tropical biodiversity. Nature, 478(7369), 378–381. 10.1038/nature10425.

Guedes, J. J. M., Moura, M. R., & Alexandre F. Diniz-Filho, J. (2023). Species out of sight: Elucidating the determinants of research effort in global reptiles. Ecography, 2023(3). 10.1111/ecog.06491.

Hortal, J., de Bello, F., Diniz-Filho, J. A. F., Lewinsohn, T. M., Lobo, J. M., & Ladle, R. J. (2015). Seven shortfalls that beset large-scale knowledge of biodiversity. Annual review of ecology, evolution, and systematics, 46, 523–549. 10.1146/annurev-ecolsys-112414-054400.

Lewin, A., Feldman, A., Bauer, A. M., Belmaker, J., Broadley, D. G., Chirio, L., Itescu, Y., LeBreton, M., Maza, E., Meirte, D., Nagy, Z. T., Novosolov, M., Roll, U., Tallowin, O., Trape, J.-F., Vidan, E. & Meiri, S. (2016). Paierns of species richness, endemism and environmental gradients of African reptiles. Journal of biogeography, 43(12), 2380–2390. 10.1111/jbi.12848.

MacKenzie, D. I., Nichols, J. D., Royle, J. A., Pollock, K. H., Bailey, L., & Hines, J. E. (2017). Occupancy estimation and modeling: inferring paierns and dynamics of species occurrence. Elsevier. 10.1016/C2012-0-01164-7.

Meiri, S., Bauer, A. M., Allison, A., Castro-Herrera, F., Chirio, L., Colli, G., Das, I., Doan, T. M., Glaw, F., Grismer, L. L., Hoogmoed, M., Kraus, F., LeBreton, M., Meirte, D., Nagy, Z. T., Nogueira, C. de C., Oliver, P., Pauwels, O. S. G., Pincheira-Donoso, D., … Roll, U. (2018). Extinct, obscure or imaginary: The lizard species with the smallest ranges. Diversity and Distributions, 24(2), 262–273. 10.1111/ddi.12678.

Myers, N., Miiermeier, R. A., Miiermeier, C. G., Da Fonseca, G. A. B., & Kent, J. (2000). Biodiversity hotspots for conservation priorities. Nature, 403(6772), 853–858. 10.1038/35002501.

Newbold, T., Oppenheimer, P., Etard, A., & Williams, J. J. (2020). Tropical and Mediterranean biodiversity is disproportionately sensitive to land-use and climate change. Nature Ecology & Evolution, 4(12), 1630–1638. 10.1038/s41559-020-01303-0.

Newbold, T., Hudson, L. N., Phillips, H. R. P., Hill, S. L. L., Contu, S., Lysenko, I., Blandon, A., Butchart, S. H. M., Booth, H. L., Day, J., De Palma, A., Harrison, M. L. K., Kirkpatrick, L., Pynegar, E., Robinson, A., Simpson, J., Mace, G. M., Scharlemann, J. P. W., & Purvis, A. (2014). A global model of the response of tropical and sub-tropical forest biodiversity to anthropogenic pressures. Proceedings of the Royal Society B: Biological Sciences, 281(1792), 20141371. 10.1098/rspb.2014.1371.

Pickersgill, M. (2007). A redefinition of Afrixalus fulvoviiatus (Cope, 1860) and Afrixalus vi`ger (Peters, 1876) (Amphibia, Anura Hyperoliidae). African Journal of Herpetology, 56(1), 23–37. 10.1080/21564574.2007.9635551.

Pauwels, O. S., Das, S., Camara, L. B., Chirio, L., Doumbia, J., D’ACOZ, C. D. U., … & Sonet, G. (2023). Rediscovery, range extension, phylogenetic relationships and updated diagnosis of the Ornate Long-tailed Lizard Latastia ornata Monard, 1940 (Squamata: Lacertidae). Zootaxa, 5296(4), 501–524. 10.11646/ZOOTAXA.5296.4.1.

Powers, R. P., & Jetz, W. (2019). Global habitat loss and extinction risk of terrestrial vertebrates under future land-use-change scenarios. Nature Climate Change, 9(4), 323–329. 10.1038/s41558-019-0406-z.

Rege, A., & Lee, J. S. H. (2023). The socio-environmental impacts of tropical crop expansion on a global scale: A case study in cashew. Biological Conservation, 280, 109961. 10.1016/j.biocon.2023.10996.

Reis-Silva F, Alves-Martins F, Martinez-Arribas J, Pizzigalli C, Seck S, Rainho A, Rocha R, Palmeirim A F (2024). Opportunistic records of amphibian and reptile across different land-use types in Guinea-Bissau, West Africa. CIBIO (Research Center in Biodiversity and Genetic Resources) Portugal. Occurrence dataset 10.15468/dwectn.

Reis-Silva F, Alves-Martins F, Martinez-Arribas J, Pizzigalli C, Seck S, Rainho A, Rocha R, Palmeirim A F (2024). Standardized survey dataset of amphibian and reptile across different land-use types in Guinea-Bissau, West Africa. CIBIO (Research Center in Biodiversity and Genetic Resources) Portugal. Sampling event dataset 10.15468/vv9xnb.

Soiomayor, M., Palmeirim, A. F., Meyer, C. F., De Lima, R. F., Rocha, R., & Rainho, A. (2024). Nature-based solutions to increase rice yield: An experimental assessment of the role of birds and bats as agricultural pest suppressors in West Africa. Agriculture, Ecosystems & Environment, 370, 109067. 10.1016/j.agee.2024.109067.

Tan, W. C., Herrel, A., & Rödder, D. (2023). A global analysis of habitat fragmentation research in reptiles and amphibians: what have we done so far?. Biodiversity and Conservation, 32(2), 439–468. 10.1007/s10531-022-02530-6.

Temudo, M. P., & Abrantes, M. B. (2013). Changing Policies, Shiving Livelihoods: The Fate of Agriculture in Guinea-Bissau: The Fate of Agriculture in Guinea-Bissau. Journal of Agrarian Change, 13(4), 571–589. 10.1111/j.1471-0366.2012.00364.x.

Temudo, M. P., & Abrantes, M. (2014). The Cashew Frontier in Guinea-Bissau, West Africa: Changing Landscapes and Livelihoods. Human Ecology, 42(2), 217–230. 10.1007/s10745-014-9641-0.

Temudo, M. P., Figueira, R., & Abrantes, M. (2015). Landscapes of bio-cultural diversity: Shiving cultivation in Guinea-Bissau, West Africa. Agroforestry Systems, 89(1), 175–191. 10.1007/s10457-014-9752-z.

Trape, J.-F., Trape, S., & Chirio, L. (2012). Lézards, crocodiles et tortues d’Afrique occidentale et du Sahara. IRD Éditions. 10.4000/books.irdedi4ons.37699.

Uetz, P., Freed, P, Aguilar, R., Reyes, F., Kudera, J. & Hošek, J. (eds.) (2023) The Reptile Database, http://www.reptile-database.org. Accessed on 01.06.2024.

WHO. (2023). https://www.who.int/news-room/fact-sheets/detail/snakebite-envenoming. Accessed on 01.06.2024.

