## Supplementary material for "Amphibian and reptile dataset across different land-use types in Guinea-Bissau, West Africa": Figures S1, S2

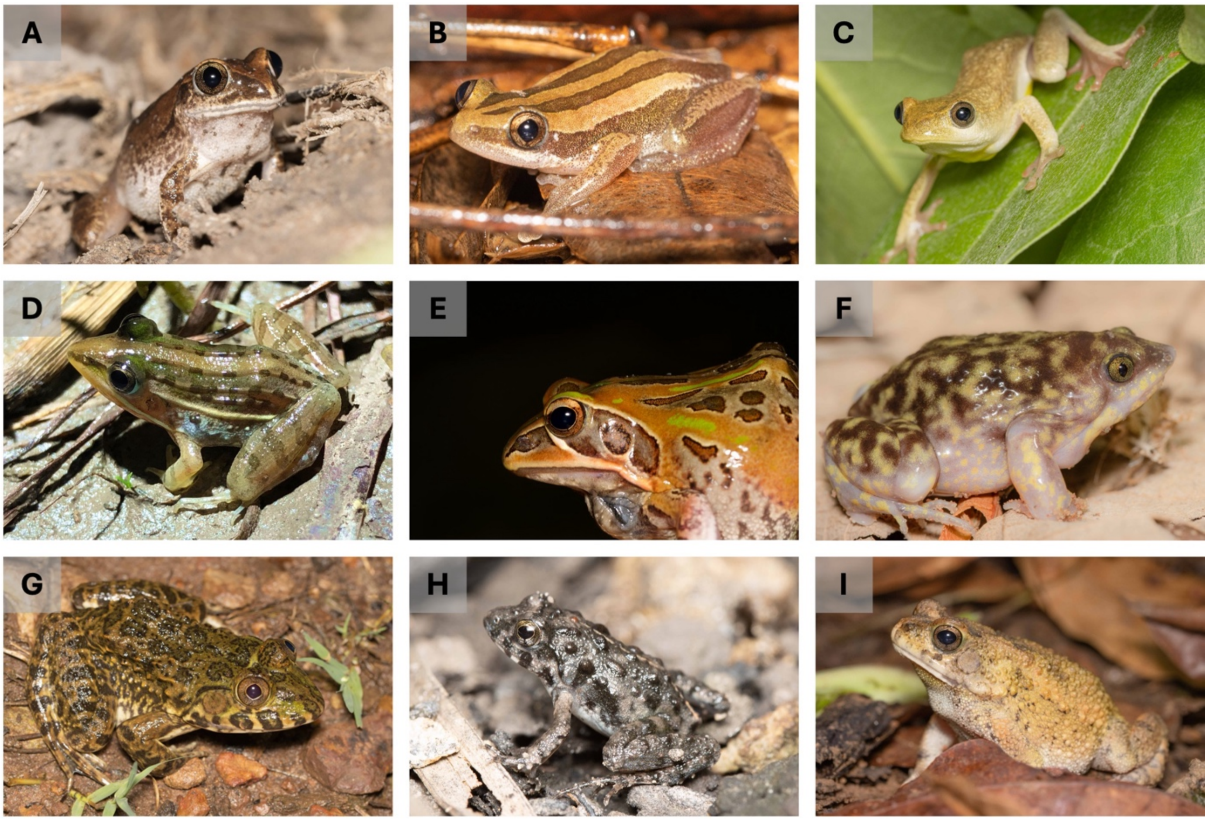


**Figure S1.** Some of the amphibians observed. A) *Leptopelis viridis*, B) *Afrixalus vittiger*, C) *Hyperolius spatzi*, D) *Ptycadena sp.*, E) *Hildebdrandtia ornata*, F) *Hemisus sp.* G) *Hoplobatrachus occipitalis*, H) *Phrynobatrachus sp.* and F) *Slerophrys sp.*. Photo credits: Francisco dos Reis-Silva.

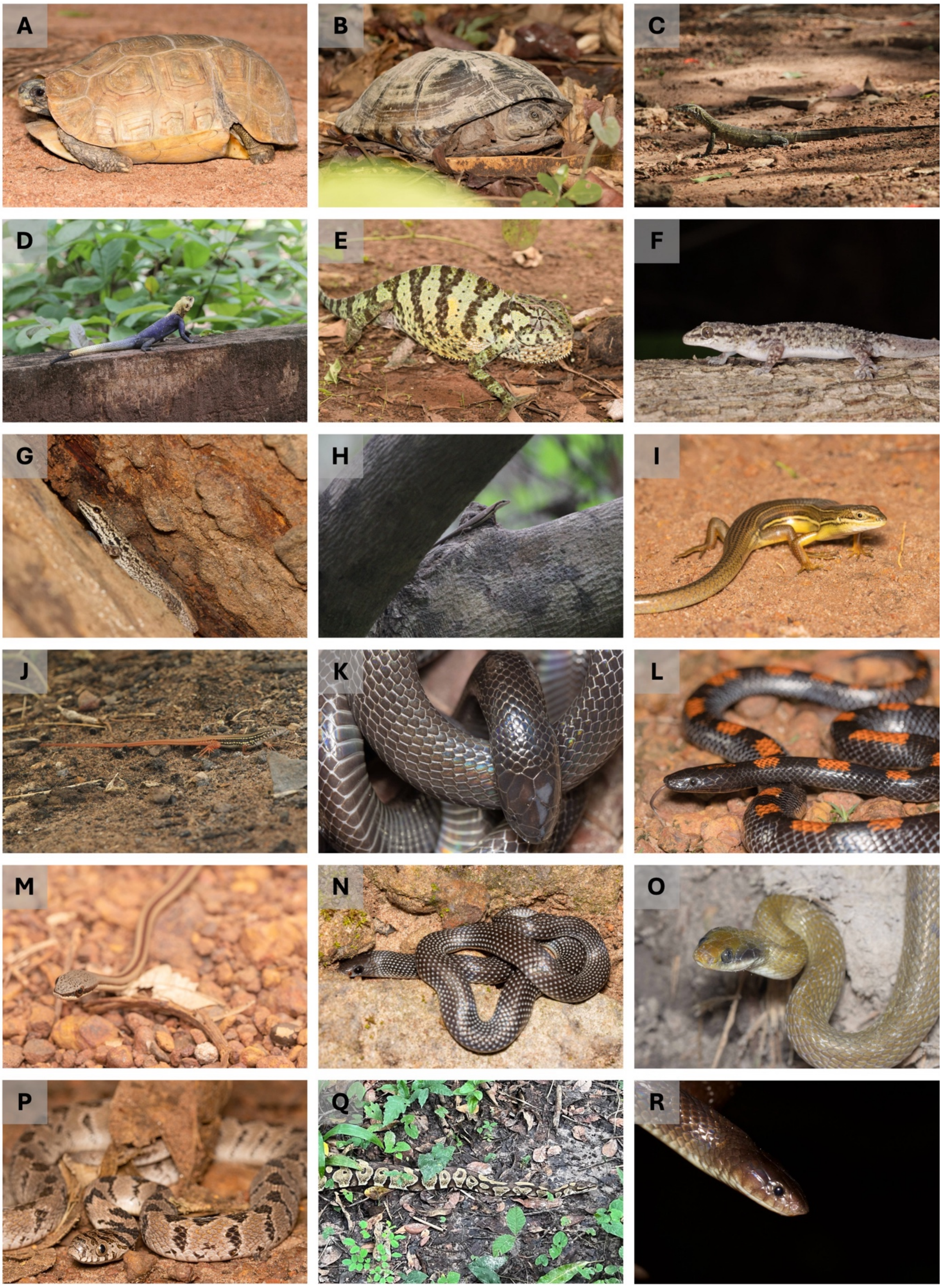


**Figure S2.** Some of the reptiles observed. A) *Kinixis belliana*, B) *Pelusios castaneus*, C) *Varanus niloticus*, D) *Agama agama*, E) *Chamaeleo gracillis*, F) *Hemidactylys angulatus*, G) *Lygodactylus gutturalis*, H) *Trachylepis affinis*, I) *Trachylepis keroanensis*, J) *Latastia ornata*, K) *Atractaspis aterrima*, L) *Lycophidion albomaculatum*, M) *Psammophis elegans*, N) *Prosymna meleagris*, O) *Crotaphopeltis*

*hotamboeia*, P) *Dasypeltis confusa*, Q) *Python regius* and R) *Elapsoidea semiannulata*. Photo credits: Francisco dos Reis-Silva (A, B, E, F, G, I, K, L, M, N, O, P), Ricardo Rocha (C, D, H, J) and Cristian Pizzigalli (Q).
